# The protein phosphatase PPKL is a key regulator of daughter parasite development in *Toxoplasma gondii*

**DOI:** 10.1101/2023.06.13.544803

**Authors:** Chunlin Yang, Emma H. Doud, Emily Sampson, Gustavo Arrizabalaga

## Abstract

Apicomplexan parasites, including *Toxoplasma gondii*, encode many plant-like proteins, which play significant roles and present attractive targets for drug development. In this study, we have characterized the plant-like protein phosphatase PPKL, which is unique to the parasite and absent in its mammalian host. We have shown that its localization changes as the parasite divides. In non-dividing parasites, it is present in the cytoplasm, nucleus, and preconoidal region. As the parasite begins division, PPKL is enriched in the preconoidal region and the cortical cytoskeleton of the nascent parasites. Later in the division, PPKL is present in the basal complex ring. Conditional knockdown of PPKL showed that it is essential for parasite propagation. Moreover, parasites lacking PPKL exhibit uncoupling of division, with normal DNA duplication but severe defects in forming daughter parasites. While PPKL depletion does not impair the duplication of centrosomes, it affects the rigidity and arrangement of the cortical microtubules. Both Co-Immunoprecipitation and proximity labeling identified the kinase DYRK1 as a potential functional partner of PPKL. Complete knockout of *DYRK1* phenocopies lack of PPKL, strongly suggesting a functional relationship between these two signaling proteins. Global phosphoproteomics analysis revealed a significant increase in phosphorylation of the microtubule-associated proteins SPM1 in PPKL-depleted parasites, suggesting PPKL regulates the cortical microtubules by mediating the phosphorylation state of SPM1. More importantly, the phosphorylation of cell cycle-associated kinase Crk1, a known regulator of daughter cell assembly, is altered in PPKL-depleted parasites. Thus, we propose that PPKL regulates daughter parasite development by influencing the Crk1-dependent signaling pathway.

**Importance:** *Toxoplasma gondii* can cause severe disease in immunocompromised or immunosuppressed patients and during congenital infections. Treating toxoplasmosis presents enormous challenges since the parasite shares many biological processes with its mammalian hosts, which results in significant side effects with current therapies. Consequently, proteins that are essential and unique to the parasite represent favorable targets for drug development. Interestingly, *Toxoplasma*, like other members of the phylum Apicomplexa, has numerous plant-like proteins, many of which play crucial roles and do not have equivalents in the mammalian host. In this study, we found that the plant-like protein phosphatase, PPKL, appears to be a key regulator of daughter parasite development. With the depletion of PPKL, the parasite shows severe defects in forming daughter parasites. This study provides novel insights into the understanding of parasite division and offers a new potential target for the development of antiparasitic drugs.

## Introduction

Apicomplexa phylum species are parasites of humans and other animals, causing various diseases such as malaria, toxoplasmosis, and cryptosporidiosis. Apicomplexa has highly specialized organelles such as micronemes, rhoptries, and polar rings, which are critical in the propagation and virulence of these parasites [1]. Among these unique organelles is the non-photosynthetic plastid known as the apicoplast, which is thought to have originated from an algal endosymbiont engulfed by a common ancestor of the current apicomplexans [2]. This secondary endosymbiotic event is also thought to have resulted in significant horizontal gene transfer from the endosymbiont to the nuclear genome during evolution [2]. As a result, apicomplexan genomes retain a multitude of plant-like genes. Many of these plant-like genes encode unique proteins essential for parasite biology, including the ApiAP2 transcription factors, which are key regulators for apicomplexan life cycle progression and differentiation [3], and the calcium-dependent protein kinases that regulate motility, invasion, and egress [4–7]. As no homologs of most of these plant-like proteins are present in mammalian cells, they serve as favorable drug targets for the development of antiparasitic drugs.

The PPP family protein phosphatase PPKL, which contains a Kelch domain at its N-terminal region, is found in land plants and green alga, and, interestingly, in alveolates, including apicomplexans [8]. All available apicomplexan genomes encode a single PPKL [8]. By contrast, *Arabidopsis* encodes four PPKLs, including brassinosteroid-insensitive1 (BRI1) suppressor (BSU1) and BSU-LIKE 1, 2, and 3 (BSL1,2,3) [9]. BSU1 is the most well-studied PPKL and is central to the brassinosteroid signaling pathway in *Arabidopsis* [10, 11]. Brassinosteroids (BRs) are essential growth-promoting hormones in plants, which are ligands of BRI1, a receptor kinase located in the plasma membrane [12, 13]. BRl1 works in conjunction with the co-receptor BRI1-associated kinase 1 (BAK1) [14]. In the absence of BRs, BRI1 is inactive and bound to the inhibitor protein BRI1 kinase inhibitor1 (BKI1) [15]. When BR binds to the extracellular domains of BRI1 and BAK1, the cytoplasmic kinase domain of BRI1 is activated and phosphorylates BKI1, leading to its dissociation [16, 17]. BRI1 is then fully activated by the formation of a heterodimeric complex with BAK1 and phosphorylation of the cytoplasmic domains of both kinases [16, 17]. Activated BRI1 initiates a signaling cascade that leads to the activation of BSU1 by phosphorylation [18, 19]. Activated BSU1 then dephosphorylates BIN2, a plant homolog of glycogen synthase kinase3 (GSK3)-like serine/threonine kinase, which leads to its degradation [19, 20]. In the absence of brassinosteroid, phosphorylated and active BIN2 phosphorylates two transcriptional factors, BZR1 (Brassinazole-resistant1) and BES1 (bri1-EMS suppressor1), leading to their retention in the cytoplasm and proteasomal degradation mediated by ubiquitination [21–23]. Thus, dephosphorylation of BIN2 by BSU1 leads to its degradation, which allows BZR1 and BES1 to act in the nucleus to activate the expression of BR-responsive genes [16, 24].

While apicomplexans do not produce BRs and lack the receptors and most of the proteins involved in brassinosteroid signaling, they express PPKLs with strong homology to BSU1. To date, the study of PPKL in apicomplexans has been limited to *Plasmodium*, where it was found to be dominantly expressed in female gametocytes and ookinetes, and deletion of PfPPKL resulted in defects in the integrity of apical structures, motility, and mosquito invasion [25]. To further explore the function of PPKL in apicomplexans, we have focused on PPKL in *Toxoplasma gondii*, an obligate intracellular parasite in the phylum Apicomplexa. *Toxoplasma* infection is prevalent in humans worldwide. Although *Toxoplasma* infection poses a minimal danger to people with healthy immune systems, it constitutes a considerable threat to immunocompromised patients and during congenital infections [26, 27]. The available drugs that can treat toxoplasmosis are very limited and have significant toxic side effects [28]. Here we show that PPKL has a highly dynamic localization pattern during parasite division and that, importantly, it is essential for parasite division. Moreover, we show that PPKL is an important part of the signaling pathways that control cell cycle, division, and cytoskeletal regulation. Therefore, this work sheds light on the unique processes by which this important pathogen divides and reveals an essential enzyme that could serve as a target for much-needed therapeutics.

## Results

### PPKL exhibits multiple cellular localizations and is associated with the progression of daughter parasite formation

PPKL in *Toxoplasma* consists of a sequence of 934 amino acids, which exhibits a structural organization like its homolog in plants and other apicomplexans, featuring six kelch motifs located in the N-terminal region, followed by the protein phosphatase domain situated in the C-terminal region (Fig. 1A). To determine where PPKL localizes within the parasite, we used a CRISPR/Cas9-based strategy to generate a strain in which the endogenous gene encoded a C-terminal triple hemagglutinin (3xHA) epitope tag. The resulting strain, Δ*ku80*:PPKL.3xHA (referred to as PPKL^HA^ hereafter), was used for immunofluorescence assays (IFA) of intracellular parasites. As transcriptomic data shows that *PPKL*’s expression varies during the cell cycle (ToxoDB), we co-stained parasites for the inner membrane complex (IMC) protein IMC3 to monitor parasite division and daughter cell formation. In non-dividing parasites, PPKL is present throughout the parasite and can be detected in both the cytoplasm and the nucleus (Fig. 1B). To confirm that PPKL is present in the nucleus, we performed cytoplasmic and nuclear fractionation of PPKL^HA^ parasites and then compared the ratio of PPKL in the nuclear fraction with that of the exclusively cytoplasmic protein eIF2α [29]. The results showed that PPKL was significantly enriched in the nuclear fraction in relation to eIF2α (Fig. S1).

**Figure 1.**
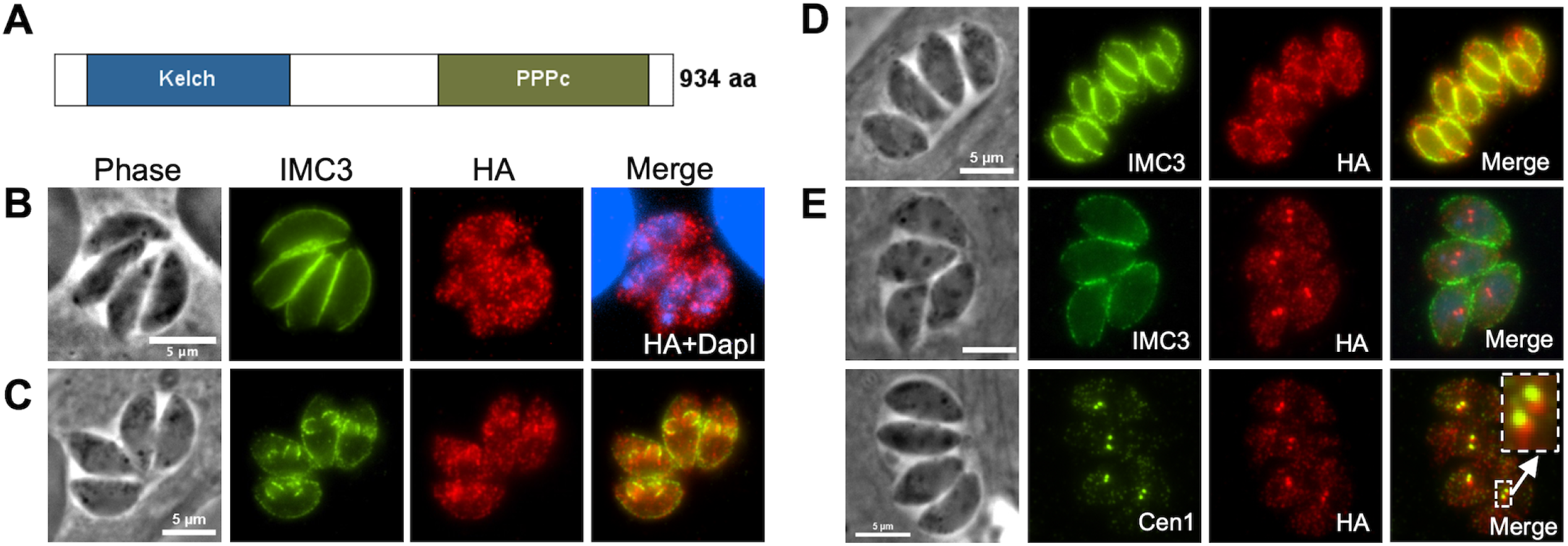
PPKL shows dynamic localization during division. A. Schematic of PPKL in *Toxoplasma*. B, C, D, and E. Intracellular parasites of the PPKL^HA^ strain were stained with anti-HA antibodies to monitor PPKL localization in non-dividing parasites (B) and in parasites in the mid (C), late (D), or early (E) stages of division. Division was monitored with antibodies against IMC3 (B-E) or Centrin1 (E). The white arrow in E indicates the area expanded in the box. Scale bar: 5 μm.

Interestingly, in dividing parasites, PPKL appears to be associated with the daughter parasites and is enriched in the cortical cytoskeleton of the daughter parasites (Fig. 1C and D). Intriguingly, we found that in a small fraction (6.5% ± 0.7%) of parasites, PPKL was heavily enriched in two distinct bright spots, which partially overlapped with the two duplicated centrosomes labeled by the anti-centrin1 antibody (Figure 1E). This localization suggests that PPKL may be present in the daughter parasite buds at a very early stage of division before the formation of IMC of the daughter cells (Fig. 1E).

For a more detailed analysis of PPKL localization, we performed Ultrastructure Expansion Microscopy (U-ExM) [30, 31]. To observe the protein and other parasite features of interest, we employed NHS-ester to stain all proteins, an acetylated-tubulin antibody to visualize microtubules, an anti-HA antibody to detect HA-tagged PPKL, and Draq5 to label DNA. As we have seen by standard IFA, analysis of U-ExM images of parasites that have expanded in volume by approximately 100-fold corroborate that PPKL is present throughout the parasite in non-dividing ones and appeared to enrich in the cortical cytoskeleton of daughter parasites (Fig. 2A, 2B). Interestingly, U-ExM allowed us to observe that PPKL is enriched at the apical end of both mother and daughter parasites in a ring-like pattern (Fig. 2A, 2B). To more clearly determine whether the PPKL ring overlapped with the apical polar ring or the preconoidal region, we performed U-ExM for extracellular parasites to observe parasites with extended conoids. In extracellular parasites with protruded conoid, the PPKL ring was on the apical end of the conoid (Fig. 2C, 2D), suggesting that PPKL is present in or near the preconoidal region. Remarkably, we observed that during the earliest stages of division, PPKL is one of the earliest components present in the daughter parasite scaffold (Fig. 2E), which only contained two rings, one labeled by PPKL, and a second one, presumably the apical polar ring, labeled by anti-acetylated tubulin antibody. In addition, we observed that in daughter parasites late in the division, PPKL was enriched in the basal complex ring (Fig. 2F), suggesting that PPKL may be involved in terminating the extension of microtubules or in the contraction of the basal complex ring. In sum, both IFA and U-ExM showed that PPKL is present in multiple cellular locations and that the dynamic localization pattern is associated with the progression of daughter parasite formation, suggesting that PPKL may play multiple roles during daughter parasite development.

**Figure 2.**
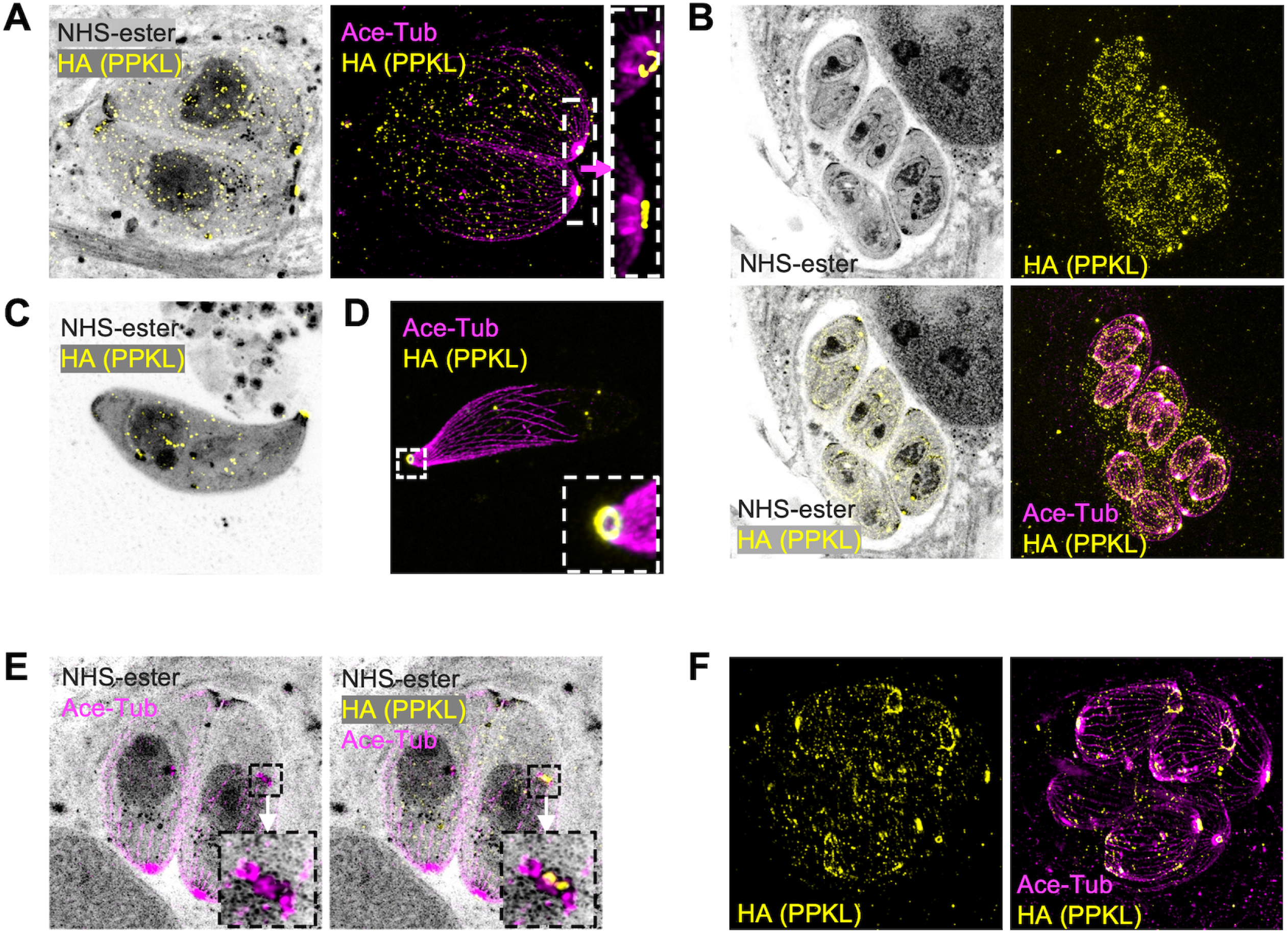
Ultrastructural expansion microscopy reveals PPKL in basal and apical structures. Intracellular and extracellular parasites were fixed with paraformaldehyde, expanded in acrylamide gels, and stained with NHS-ester, anti-HA, and anti-acetylated tubulin. Images were captured by LSM 900 with Airyscan. A. Image of intracellular non-dividing parasites. The white box frames the region expanded to the right of the arrow. B. Images of intracellular dividing parasites. C, D. Images of extracellular parasites. The white framed zone in D is zoomed in and shown in the lower right corner. E. Image of intracellular parasites. The parasite on the right has started daughter parasite assembly, as shown by duplicated centrosomes, preconoidal regions, and the apical polar rings, which are framed in a black box. The box is enlarged to the left showing the acetylated tubulin signal, and to the right showing both anti-acetylated tubulin and anti-HA. F. Image shows intracellular parasites in a late stage of division.

### Depletion of PPKL in parasites leads to disruption of parasite division

Through a *Toxoplasma* genome-wide CRISPR screen, PPKL was assigned a fitness score of -5.02 [32], suggesting that it is essential for parasite survival and, therefore, that knockout of the *PPKL* gene is not likely possible. Accordingly, to investigate its function, we generated a conditional knockdown strain by using the auxin-induced degradation (AID) system [33]. For this purpose, we inserted an AID-3xHA prior to the stop codon of the endogenous *PPKL* gene using CRISPR-mediated gene editing. Western blot of the resulting PPKL^AID-HA^ strain showed that treatment with auxin significantly reduced the amount of PPKL within half an hour and almost completely depleted PPKL within one hour (Fig. 3A). To determine if PPKL is required for *Toxoplasma* propagation, we performed plaque assays of both the parental and PPKL^AID-HA^ strains with and without auxin. Consistent with the low fitness score, PPKL^AID-HA^ parasites treated with auxin failed to form any plaques (Fig. 3B and C), indicating that PPKL is essential for parasite propagation. In addition, we observed that untreated PPKL^AID-HA^ parasites formed fewer and smaller plaques than the parental strain (Fig. 3B and C), which may be due to the significantly lower expression of PPKL-AID in relation to the parental strain (Fig. S2) or perhaps because the fusion of AID to PPKL may slightly impair its function.

**Figure 3.**
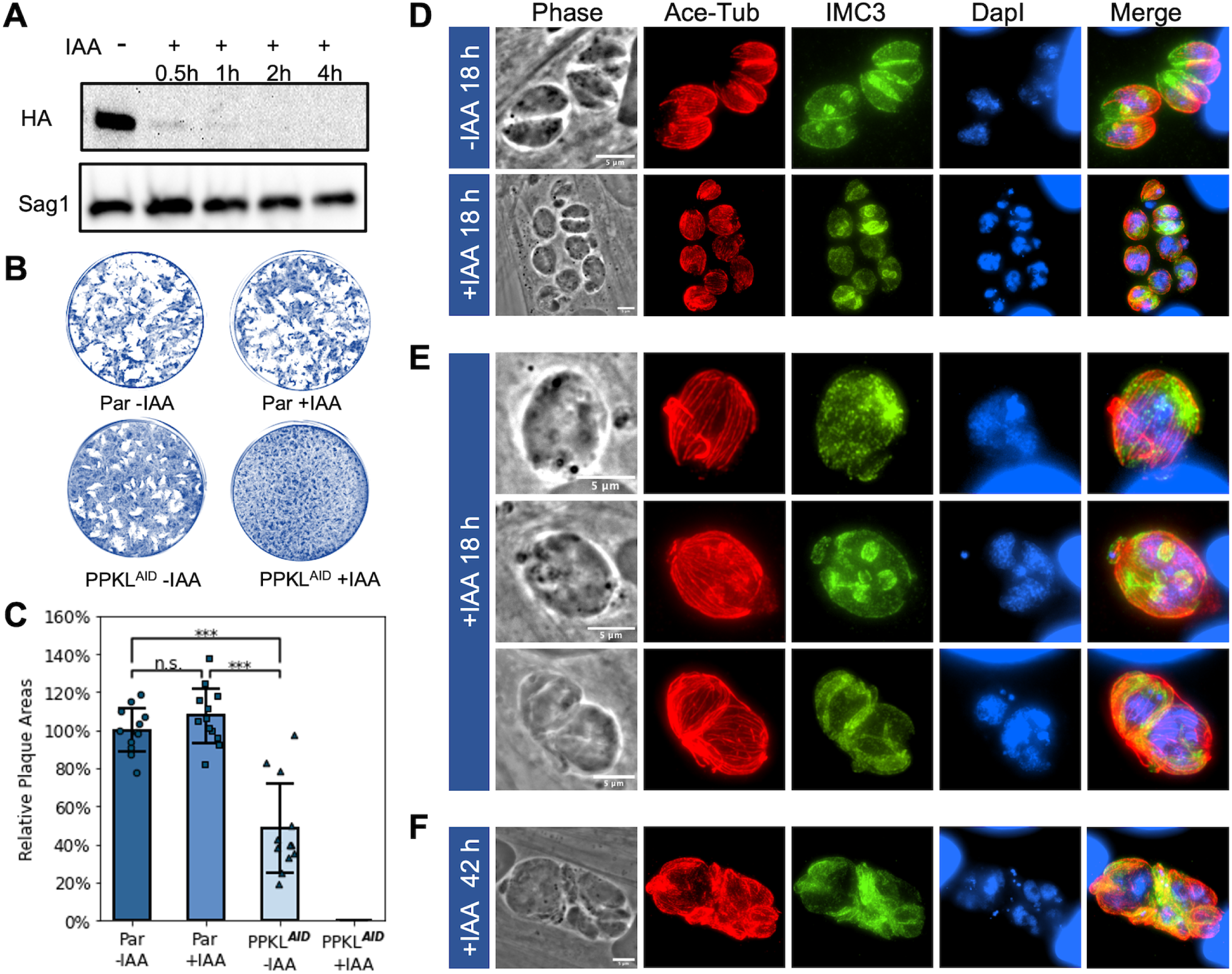
Depletion of PPKL leads to disruption of parasite division. A. Shown is representative Western blot of protein extract of the AID-tagged PPKL strain (PPKL^AID^) treated with auxin (IAA) for times indicated and probed for the HA epitope tag. The protein Sag1 was used as a loading control. B and C. Parasites from the parental (par) or PPKL^AID^ were grown for 6 days in the absence or presence of auxin (IAA) and allowed to form plaques. B shows representative plaque assays. The graph in C is the average plaque area formed by the two strains, based on data collected from four biological replicates, with three experimental replicates for each. For each biological replicate, the data was normalized to the average plaque area of the parental strain without IAA treatment. The error bars represent standard deviations. ***, P<0.001; n.s., no significance (Student’s t-test, two tails, unequal variance). D to F. PPKL^AID^ parasites were grown with/without auxin for 18 hours (D and E) or 42 hours (F) and analyzed by IFA. The cultures were stained with anti-acetylated tubulin, anti-IMC3, and DapI. Scale bar: 5 μm.

To determine the cellular consequences of PPKL depletion, we performed IFA to observe the morphology of PPKL^AID-HA^ parasites treated with auxin. Briefly, we added auxin to the culture 2 hours after infection and allowed parasites to grow overnight (∼18h). IFAs were performed using antibodies against acetylated tubulin and IMC3 to monitor the formation of daughter cells. Strikingly, in the auxin-treated PPKL^AID-HA^ cultures, most vacuoles contained only one extremely swollen parasite, whereas untreated vacuoles mostly had 4 to 8 parasites (Fig. 3D). Interestingly, these swollen parasites contained a large amount of DNA, far exceeding the amount of a normal single parasite (Fig. 3D and E). However, most (71% ± 6%) of these swollen parasites did not have any daughter parasites (Fig. 3E, top row), and only a few (10% ± 2%) contained a varied number of seemingly abnormal daughter parasites (Fig. 3E, middle row). Moreover, there were some (18% ± 4%) vacuoles containing two expanded parasites, similarly with an increased amount of DNA (Fig. 3E, bottom row). When we extended the treatment of auxin to two days (∼42h), the vacuoles became more disorganized, had more abnormal parasites, and swollen parasites had a further accumulation of DNA (Fig. 3F), implying that division was progressing, albeit abnormally. In summary, PPKL-depleted parasites have deficiencies in daughter parasite formation. Still, their ability to replicate DNA appears relatively unimpaired, resulting in severe division uncoupling.

### Depletion of PPKL does not affect the replication of centrosomes

During endodyogeny, the two daughter parasite scaffolds are assembled near the two centrosomes in a one-to-one correspondence [34]. Centrosome duplication has been shown to occur before the formation of the daughter parasites, and it is needed for the division to initiate [35, 36]. Based on the significant defects in parasite division in the PPKL mutant, we monitored centrosome duplication upon PPKL depletion. As expected, untreated dividing parasites had exactly two centrosomes in each mother cell (Fig. 4A). Similarly, parasites treated with auxin overnight had at least two centrosomes (Fig. 4A), regardless of whether they contained daughter parasites or not, implying that centrosome duplication was still taking place. Interestingly, we often observe treated parasites with more than two centrosomes (Fig. 4A). It should be considered that, under normal conditions, parasites that have been cultured overnight would have undergone two to three divisions. Thus, the presence of more than two centrosomes within one parasite is consistent with our observation that PPKL depletion leads to division (i.e., centrosome and DNA duplication) and daughter parasite formation being uncoupled.

**Figure 4.**
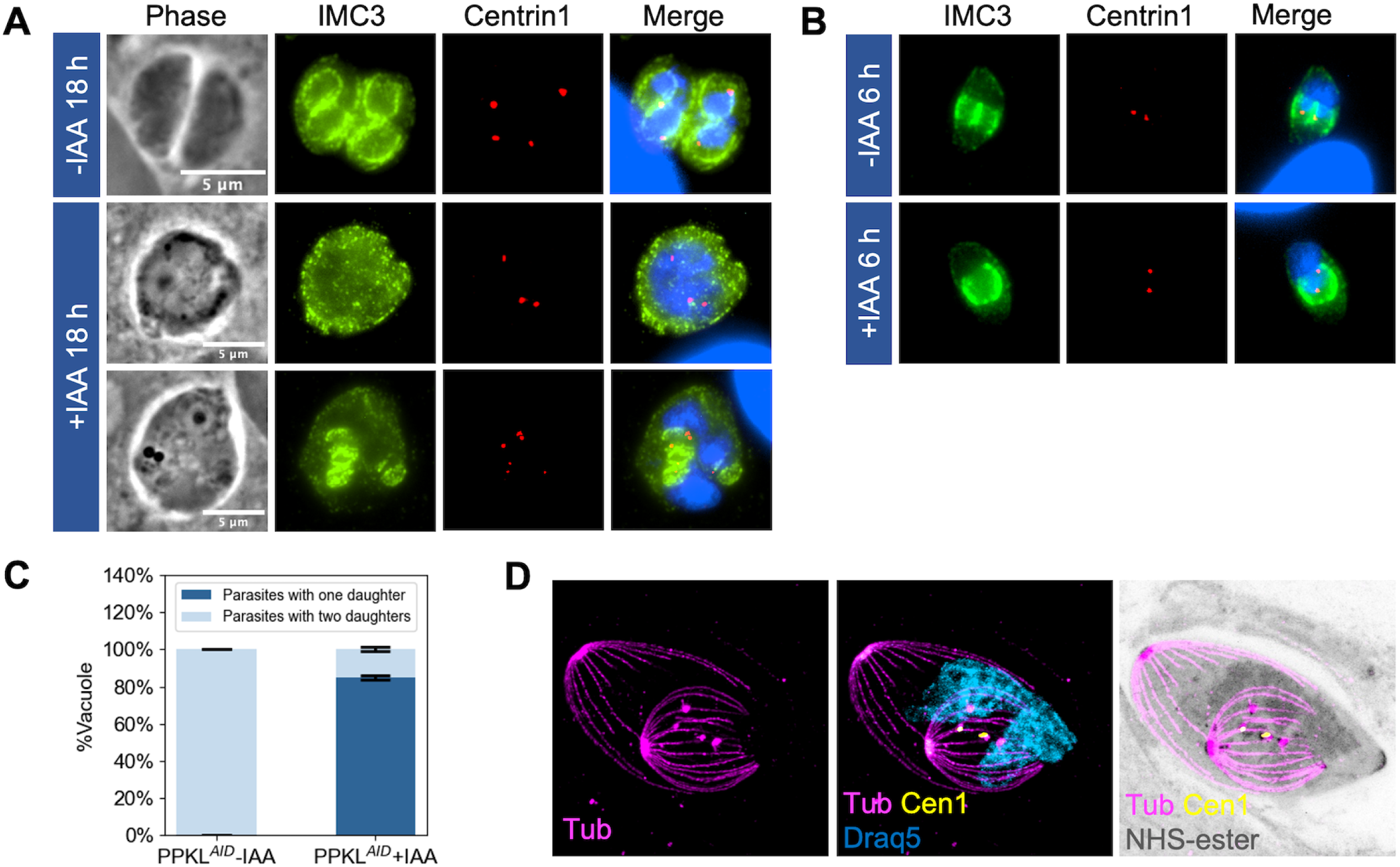
Depletion of PPKL does not affect the replication of centrosomes. Centrosome duplication of PPKL^AID^ grown with and without auxin was monitored by IFA using antibodies against centrin1. A. IFA of parasites grown with/without IAA for 18 hours and stained with anti-IMC3, anti-Centrin1 antibodies, and DapI. B. IFA of PPKL^AID^ parasites grown with/without IAA for 6 hours. C. quantification of dividing PPKL^AID^ parasites treated with/without IAA for 6 hours with one or two daughters. Three biological replicates were performed, with 50 parasites counted for each replicate. The error bars represent standard deviations. D. U-ExM of one daughter containing PPKL^AID^ parasite treated with IAA for 6 hours and stained with anti-acetylated tubulin, anti-Centrin1, NHS-ester, and Draq5. Scale bar: 5 μm.

To further examine centrosome duplication in PPKL-depleted parasites, we added auxin right after infection and allowed the parasites to grow for six hours, which allowed us to monitor the first division cycle of the parasites after PPKL depletion. IFA showed that few parasites had detectable daughter parasites for both the control and auxin treated. Interestingly, among the parasites that were dividing, 85%±1.5% of those from auxin-treated parasites had only one daughter parasite (Fig. 4B and C). 95%±1.5% of the parasites containing only one daughter had two centrosomes. We also observed that in the treated parasites, the single-daughter parasite appeared to be morphologically unusual and larger than normal. We explored this phenotype using U-ExM and observed that the microtubule system of the single daughter parasite appeared to be normal, except that the gaps between the cortical microtubules were wider, which correlates with the larger appearance of the parasites (Fig. 4D). As shown in Fig. 4D, we also observed parasites in which there were two centrosomes and two spindles, with only one of the spindles associated with the DNA. Taken together, our results suggest that although division is impaired upon depletion of PPKL, centrosome duplication remains unaffected.

### Depletion of PPKL reduces the rigidity of microtubules

A previous study has shown that the knockout of the PPKL gene in *Plasmodium* led to apical microtubule disorganization and dissociation from the IMC [25]. Interestingly, PPKL also appears to be required to maintain the order and/or rigidity of microtubules in *Toxoplasma,* as observed through the disruption of the compact structure of the cortical cytoskeleton in the knockdown strain. To confirm this observation, we treated PPKL^AID-HA^ parasites that had undergone two to three rounds of division with auxin for six hours and isolated their cortical cytoskeleton using sodium deoxycholate. As expected, the cortical cytoskeleton from the control parasites appeared more compact with few fragmented microtubules (Fig. 5A). By contrast, the cortical cytoskeleton from auxin-treated parasites appeared more disordered with five times more fragmented microtubules than the control (Fig. 5A, B), suggesting that PPKL is involved in the regulation of the rigidity and compact structure of the cortical microtubule system.

**Figure 5.**
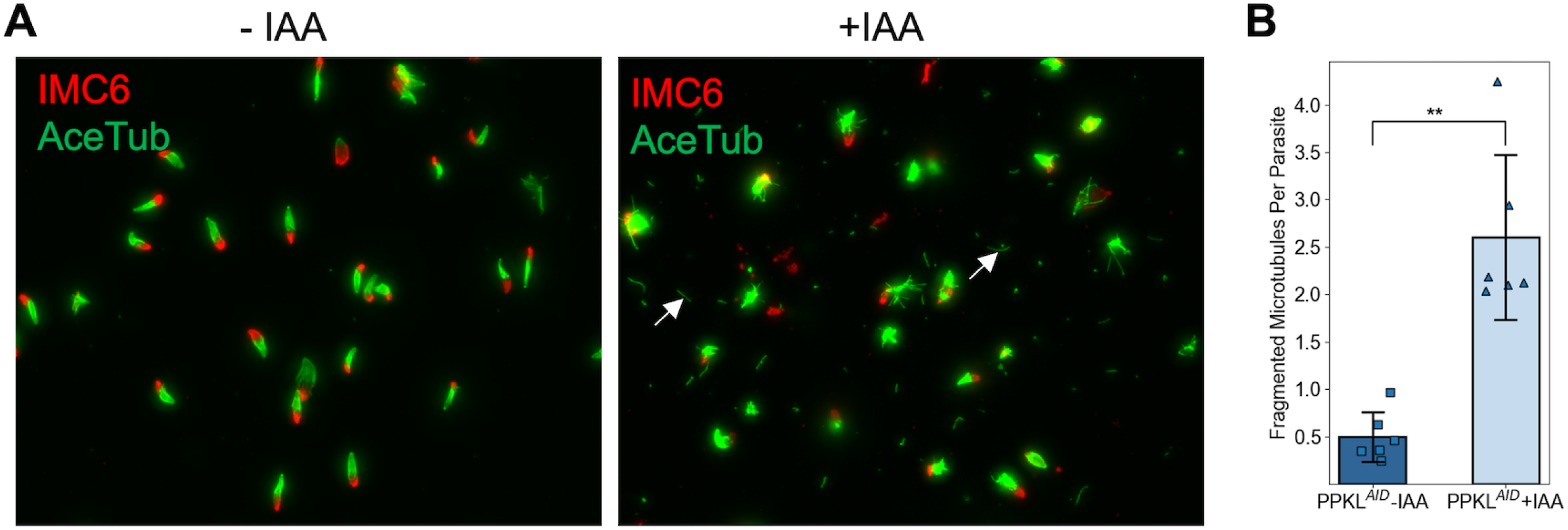
Depletion of PPKL reduces the rigidity of microtubules. The cortical cytoskeleton was extracted from PPKL^AID^ parasites treated with IAA or ethanol for 6 hours. A. IFA of the extracted cortical cytoskeleton used anti-acetylated tubulin and anti-IMC6 antibodies to monitor the microtubules and IMC, respectively. Examples of fragmented microtubule are indicated by white arrows. B. Quantification of fragmented microtubules normalized to the number of cortical cytoskeletons. The bars indicate the average number of fragment microtubules from each cortical cytoskeleton. The error bars represent standard deviations. Three biological replicates, each consisting of two experimental replicates, were performed. For each experimental replicate, 10 random fields of view were selected to count the number of cortical cytoskeleton and fragmented microtubules. **, P<0.01 (Student’s t-test, two tails, unequal variance).

### IMC29 and DYRK1 are likely PPKL functional partners

To determine the molecular mechanisms responsible for the PPKL knockdown phenotypes, we set out to identify functional partners and putative substrates of PPKL. We first performed standard co-immunoprecipitation (Co-IP) using anti-HA conjugated magnetic beads to precipitate HA-tagged PPKL from the PPKL^HA^ parasites. However, this approach failed to identify PPKL interactors, suggesting that either PPKL works on its own or that it interacts with other proteins and substrates transiently or with low affinity. Accordingly, we used disuccinimidyl sulfoxide (DSSO) to crosslink the protein samples before performing Co-IP. The results of the tandem mass spectrometry (MS/MS) analysis indicated that two proteins significantly co-precipitated with PPKL: IMC29 (TGGT1_243200) and DYRK1 (TGGT1_204280) (Supplementary Dataset 1). The inner membrane IMC29 has been identified as a crucial component of the early daughter buds with a substantial contribution to parasite division [26]. DYRK1 is a cell-cycle-related protein kinase of unknown function.

As a complementary approach to the Co-IP, we used the TurboID proximity labeling method [37], which can identify neighboring and interacting proteins, including substrates. To this end, we endogenously fused the engineered biotin ligase TurboID and a 3xHA epitope tag to the C-terminus of PPKL. IFA of the obtained parasite strain showed that PPKL^TurboID.HA^ has the same localization pattern as PPKL^HA^ (Fig. S3A). We then confirmed that the fusion was active by incubating the PPKL^TurboID.HA^ parasites with D-biotin and performed western blot, which showed a significant increase in biotinylated proteins compared to the same parasite strain without biotin treatment (Fig. S3B). To identify interacting and neighboring proteins, we incubated PPKL^TurboID.HA,^ or parental strain parasites with D-biotin and isolated biotinylated proteins for MS/MS for two experimental replicates. We applied the following criteria to the resulting list of putative interactors: 1) a total of 10 or more between the two replicates, and 2) an experiment to control fold change equal to or larger than 3.5. In this manner, we obtained a list of 81 putative PPKL neighboring and/or interacting proteins (Table 1 and Supplementary Dataset 2). To further analyze this list of proteins, we investigated their potential molecular functions based on functional annotations on ToxoDB. Out of the 81 proteins, 19 are related to the cortical cytoskeleton system, including the IMC, the apical complex, the basal complex, and microtubules; 7 are potentially related to vesicle transport; 5 are potentially related to RNA splicing (Table 1). Moreover, there are three protein kinases (TGGT1_231070, DYRK1, and SRPK1) and one protein phosphatase, PPM2A. Notably, both IMC29 and DYRK1, which were identified as putative interactors via crosslinking and IP, were identified with the TurboID approach, suggesting that they are highly likely functional partners of PPKL.

**Table 1.**
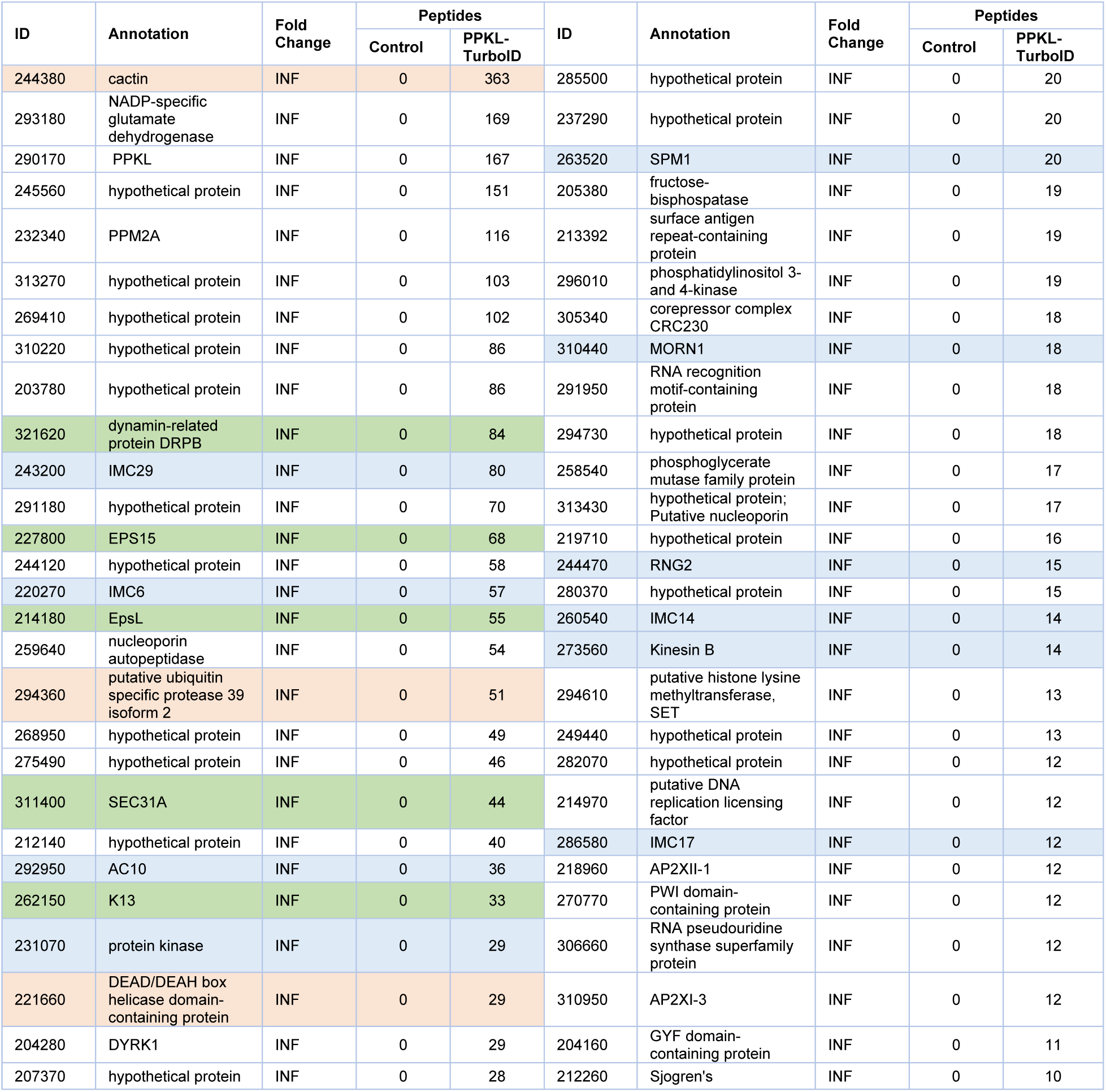
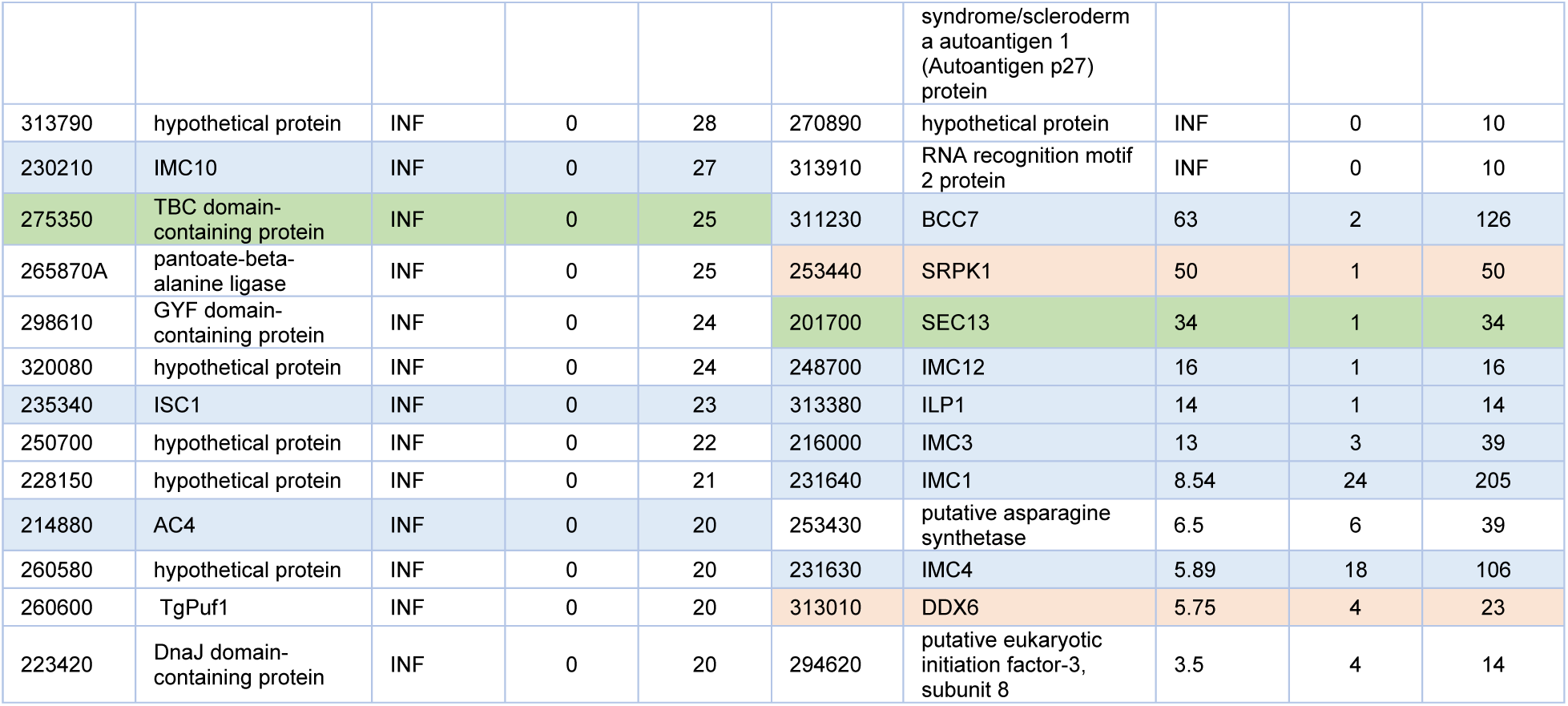
Putative PPKL neighboring proteins. Listed are proteins identified by TurboID that met the following criteria: 1) identified in both replicates; 2) a total of 10 or more peptides identified between the two replicates; 3) fold change between experiment and control equal to or larger than 3.5. Proteins related to the IMC, Apical, Basal complex, and microtubules are highlighted with light blue; proteins related to vesicle transport were highlighted with light green; and proteins related to RNA splicing are highlighted with light orange. The number of peptides listed is the total between the two replicates.

### DYRK1 plays an important role in the *Toxoplasma* division

Based on the division phenotype of PPKL-depleted parasites, it is likely that PPKL participates in a signaling pathway that regulates parasite division. As signaling pathways often involve a chain of kinases and phosphatases, we focused on characterizing the function of DYRK1, which appears to be a putative PPKL interactor. Dual-specificity tyrosine-regulated kinase (DYRK) is a member of the CMGC group of kinases [38], a large and conserved family of kinases that play key roles in cell cycle regulation and many important signaling pathways [39]. *Toxoplasma* encodes two DYRKs in its genome, and phylogenetic analysis (Fig. 6A), including human and Arabidopsis homologs, showed that DYRK1 was exclusively clustered with the plant homologs, while DYRK2 (TGGT1_283480) was clustered with human homologs. Thus, both PPKL and DYRK1 are closer in homology to proteins from plants. To identify the localization of DYRK1, we endogenously fused a 3xMyc ectopic tag at its C-terminus with CRISPR/Cas9 mediated gene editing. IFA showed that in non-dividing parasites, DYRK1 localizes to the nucleus (Fig. 6B top row); while in parasites undergoing division, it exclusively localizes to the IMCs of daughter parasite buds (Fig. 6B bottom row). The dynamic localization of DYRK1 suggests that this kinase might have multiple roles throughout the division cycle of the parasite.

**Figure 6.**
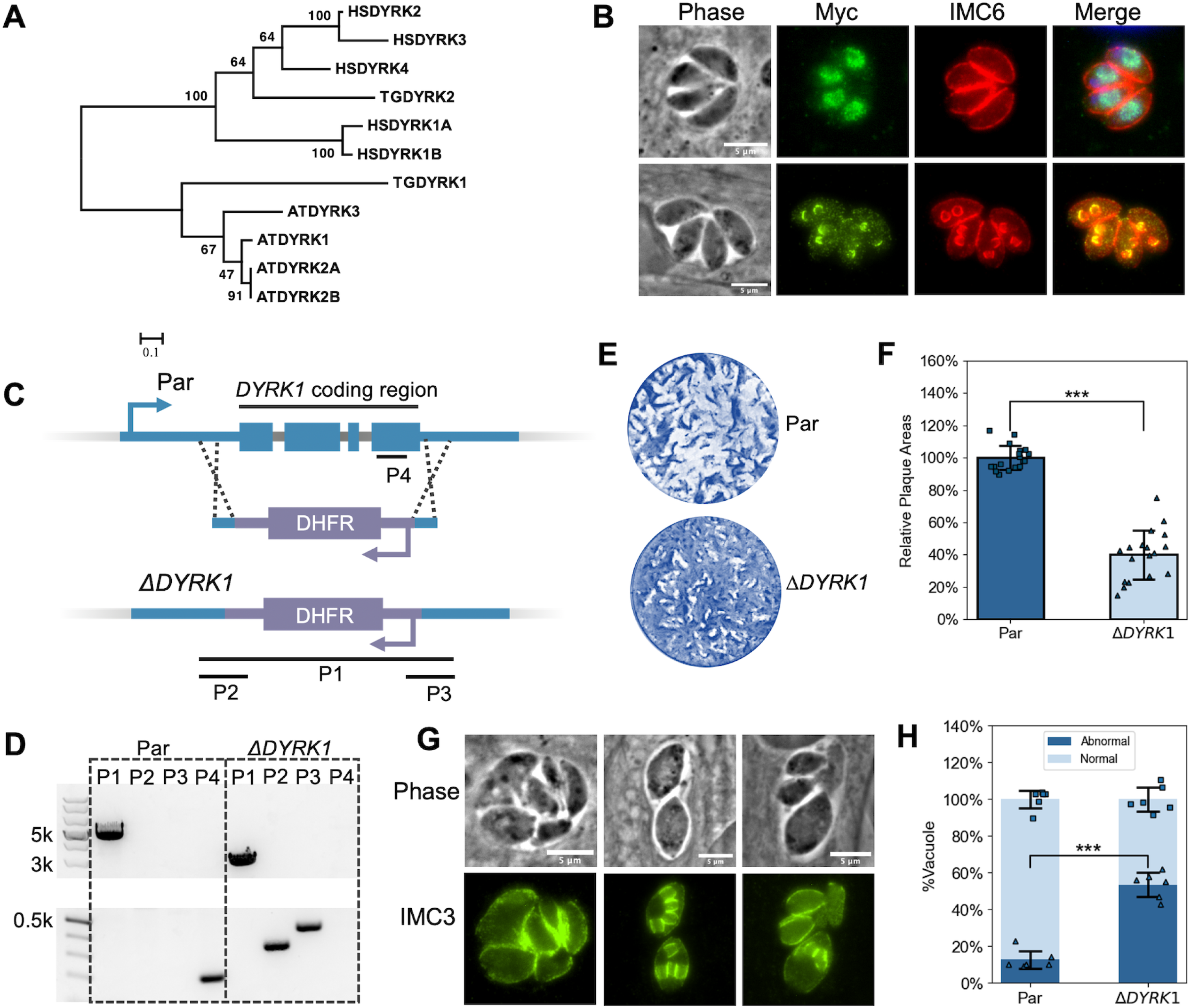
DYRK1 is a plant-like kinase and plays an important role in parasite division. A. Phylogenetic analysis of DYRK sequences from humans (HSDYRK1A, 1B, 2, 3, and 4), *Arabidopsis* (ATDYRK1, 2A, 2B, and 3), and *Toxoplasma* (TGDYRK1 and 2). DYRKs from *Toxoplasma* are highlighted. Alignment and tree construction details and accession numbers are listed in the Methods session. B. IFA of parasites expressing Myc-tagged DYRK1 using anti-Myc and anti-IMC6 antibodies. The upper panel shows non-dividing parasites, while the lower panel shows dividing ones. Scale bar: 5 μm. C. The diagram depicts the CRISPR/Cas9-mediated strategy used to disrupt the *DYRK1* gene. P1, 2, 3, and 4 are the three amplicons used to confirm the integration of the DHFR cassette between the Cas9 cutting sites and the deletion of the *DYRK1* gene. D. Agarose gel of PCR products amplified from the genomic DNA extracted from Δ*DYRK1* parasites using the four sets of primers indicated in C. The P1 amplicon has a length of 2928 bp in the Δ*DYRK1* genome and 4787 bp in the parental genome. The P2, P3, and P4 amplicons have a length of 339 bp, 431 bp, and 196 bp, respectively. E and F. Δ*DYRK1* and parental strain parasites were allowed to form plaques in culture for six days. A representative plaque assay (E) and the quantification are shown (F). The bars represent the relative average plaque areas. Three biological replicates were performed, and six experimental replicates were included each time. For each biological replicate, the data was normalized to the average plaque area of the parental. G. IFA images of Δ*DYRK1* parasites that show abnormal division. H. Quantification of the ratio of vacuoles containing parasites displaying abnormal division or morphologies. *** P<0.001 (Student’s t-test, two tails, unequal variance).

As a potential functional partner of PPKL, we sought to investigate whether the absence of DYRK1 manifests phenotypic similarities to the PPKL knockdown. For this purpose, the *DYRK1* gene was disrupted through the replacement of the coding region with a DHFR expression cassette in the parental Δ*ku80* parasites by using the CRISPR/Cas9 system (Fig. 6C). The disruption of the *DYRK1* locus in the resulting Δ*DYRK1* strain was confirmed by PCR with four sets of primers (Fig. 6D). Plaque assay showed that Δ*DYRK1* parasites are less efficient at forming plaques as compared to the parental strain (40 ± 15 % plaquing efficiency relative to parental, Fig. 6E, F). While these data suggest that DYRK1 is not essential, it appears to play an important role in parasite propagation. Importantly, IFA revealed that 53.5% ± 6.5% of vacuoles formed by Δ*DYRK1* parasites displayed abnormal parasite division (Fig. 6G, H), characterized by predominantly unsynchronized division, resulting in disorganized parasites within the vacuole and even atypical morphologies for some parasites, such as swollen parasites similar to what is observed upon PPKL depletion. The observed division defect, stemming from the deletion of DYRK1, strongly indicates the significant involvement of DYRK1 in regulating parasite division and a potential functional relationship between PPKL and DYRK1.

### PPKL influences the phosphorylation state of regulators of microtubule rigidity and cell division

To further explore the molecular mechanisms leading to the division phenotypes of PPKL-depleted parasites, we used tandem mass tag (TMT) quantitative mass spectrometry to compare the phosphoproteomes of PPKL^AID^ parasites treated with/without auxin. Briefly, PPKL^AID-HA^ parasites were allowed to grow for 18 hours before adding either auxin or ethanol (vehicle control). After one, three, and six hours of treatment, cultures were harvested, and the parasites were released by syringe lysis, and samples were prepared for quantitative mass-spectrometry. At the 6-hour timepoint, we identified 486 phosphopeptides from 313 proteins that were more than 2-fold abundant in the auxin-treated parasites (Fig. 7A, Supplementary dataset 3). We also identified 425 phosphopeptides from 255 proteins that were more than 2-fold less abundant upon PPKL depletion (Fig. 7A, Supplementary dataset 3). Interestingly, 86 proteins had both over and under-phosphorylated peptides. Thus, we identified a total of 482 proteins whose phosphorylation state was PPKL-dependent. Interestingly, 24 of the 81 proteins identified in the TurboID assay were among these 482 proteins (Supplementary Dataset 3), strongly validating the reliability of the phosphoproteome and interactome data. However, DYRK1 was not among these 24 proteins, and it had a 1.22-fold increase in phosphorylation of one residue and a 1.16-fold decrease in phosphorylation of another upon depletion of PPKL.

**Figure 7.**
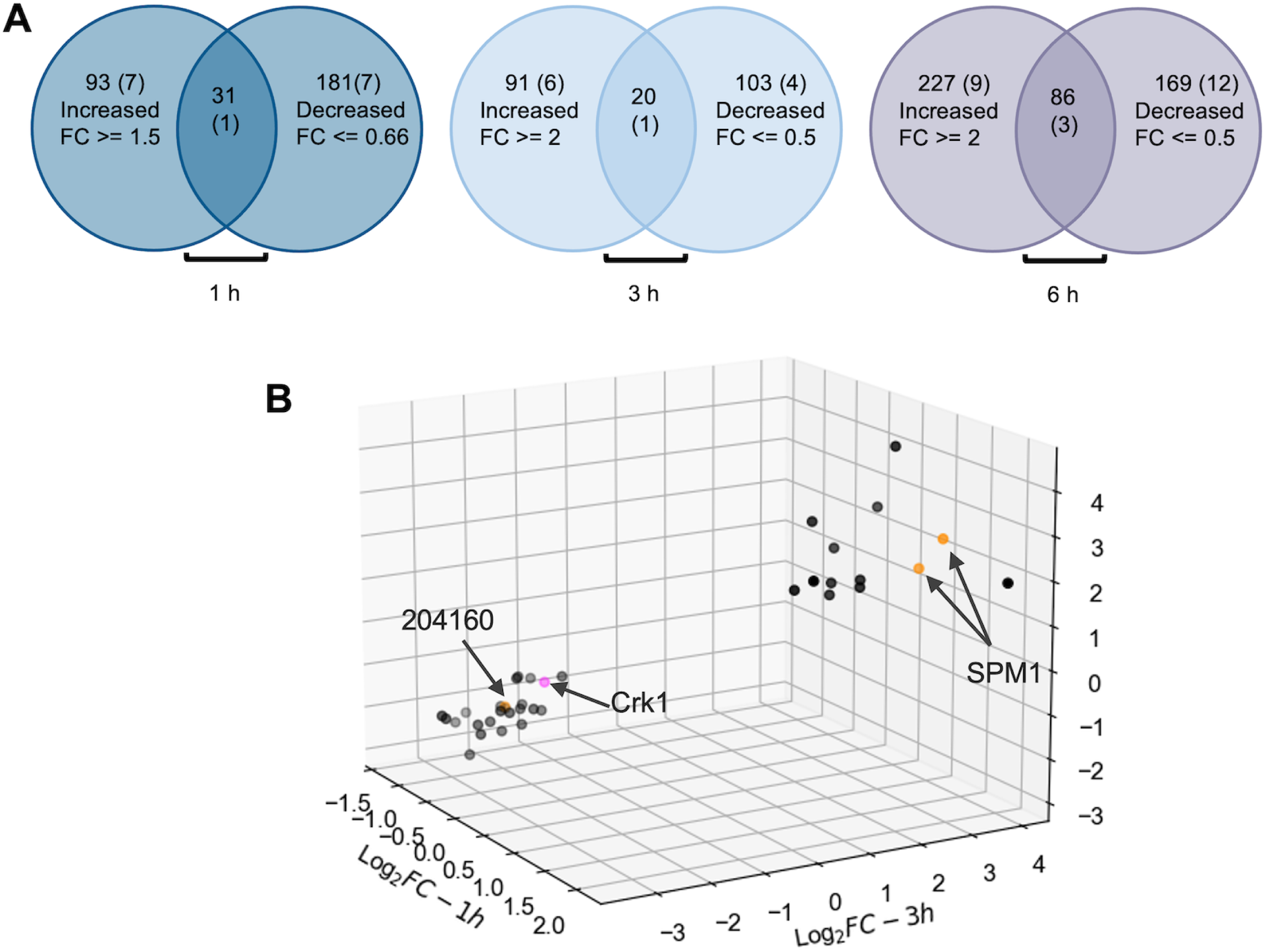
Phosphoproteome analysis showed that depletion of PPKL results in increased phosphorylation of SPM1 and decreased phosphorylation of Crk1. A. Venn diagrams showing the number of proteins with phospho-peptides that were more or less abundant in PPKL^AID^ parasites treated with auxin for 1, 3 and 6 hours. The overlap indicates the number of proteins that had phosphopeptides that increased and decreased in abundance upon PPKL depletion. The number of proteins that were also identified as PPKL neighboring proteins in each category is shown in parentheses. The fold change cutoffs are as follows: for 1 h timepoint sample set, phospho-peptides with FC >=1.5 and FC <= 0.66, for 3 h and 6 h timepoints sample sets, phospho-peptides with FC >= 2 and FC <= 0.5. B. 3D Scatter plot shows 35 phospho-peptides that are shared by all the three timepoint sample sets after filtering with the cutoffs described above. Each dot in the 3D Scatter plot represents one phosphopeptide. The dots shown in orange are peptides from PPKL neighboring proteins identified by TurboID analysis. The dot shown in pink represents the peptide from Crk1. X-axis: the log2 fold change of 1 h samples. Y-axis: the log2 fold change of 3 h samples. Z-axis: the log2 fold change of 6 h samples.

As expected, the results from the parasites treated for either one or three hours revealed a more limited number of peptides exhibiting PPKL-dependent phosphorylation status, especially for the 1-hour treated samples (Supplementary Dataset 4). Therefore, when filtering the quantitative proteomic data, we followed the 2-fold change as the cutoff for 3-hour timepoint samples but reduced the fold change from 2 to 1.5 for the 1-hour timepoint samples to prevent missing important information. Thus, we identified 204 proteins (over: 124 peptides from 101 proteins; under: 153 peptides from 123 proteins; both: 20 proteins) that exhibited PPKL-dependent phosphorylation state in the 3-hour timepoint samples, and 304 proteins (over: 142 peptides from 124 proteins, under: 278 peptides from 212 proteins; both: 32 proteins) in the 1-hour timepoint samples (Fig. 7A, Supplemental Dataset 4). When combining the data to identify phosphopeptides shared by all three time points samples, we found 13 phosphopeptides from 11 proteins were over-phosphorylated, and 22 phosphopeptides from 19 proteins were under-phosphorylated (Table 2, Fig. 7B). One of these proteins, TGGT1_214270, had both over-and under-phosphorylated peptides identified. Thus, we obtained a list of 29 proteins whose phosphorylation was affected by lack of PPKL at all three time points of auxin treatment (Table 2).

**Table 2.**
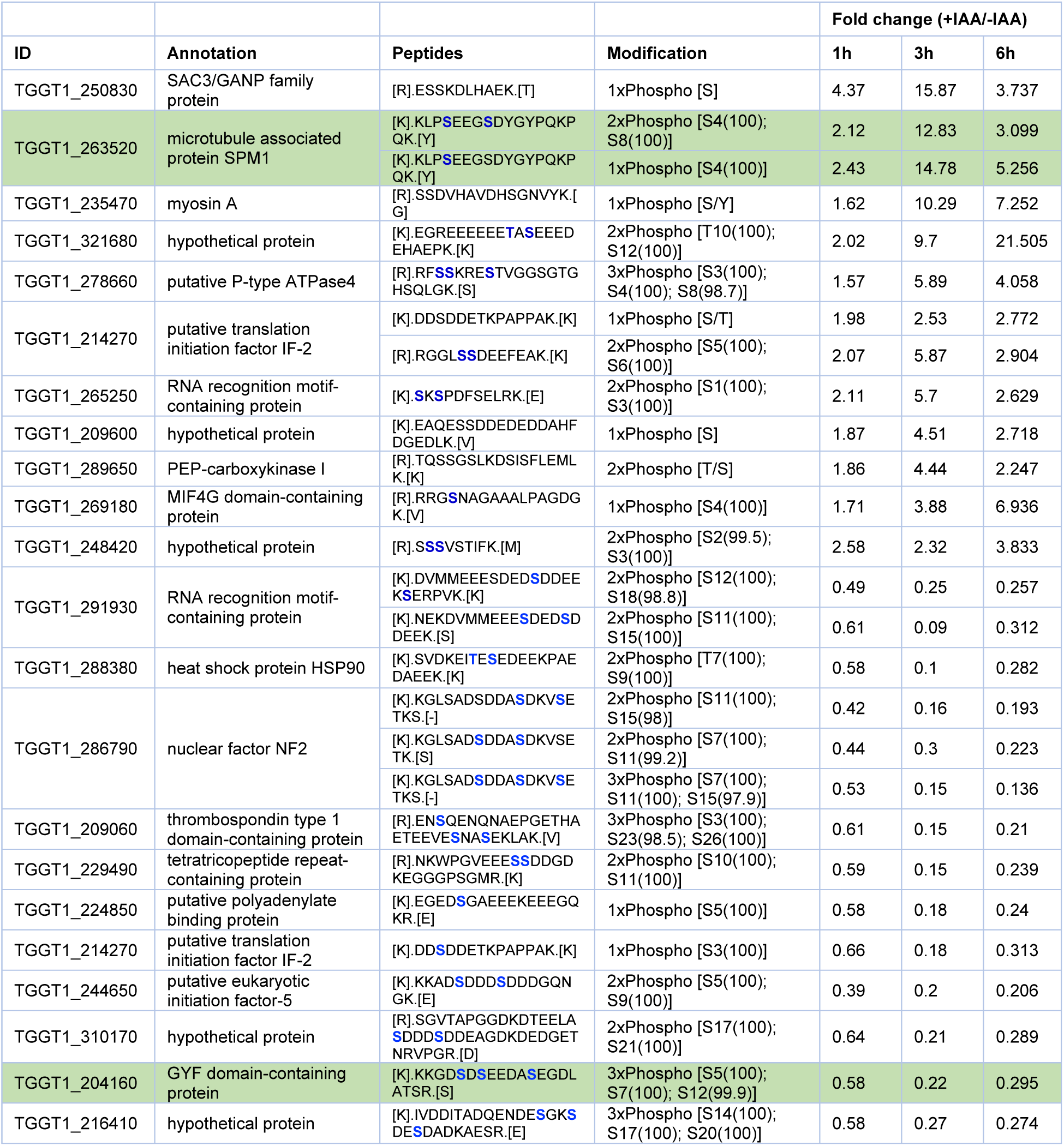
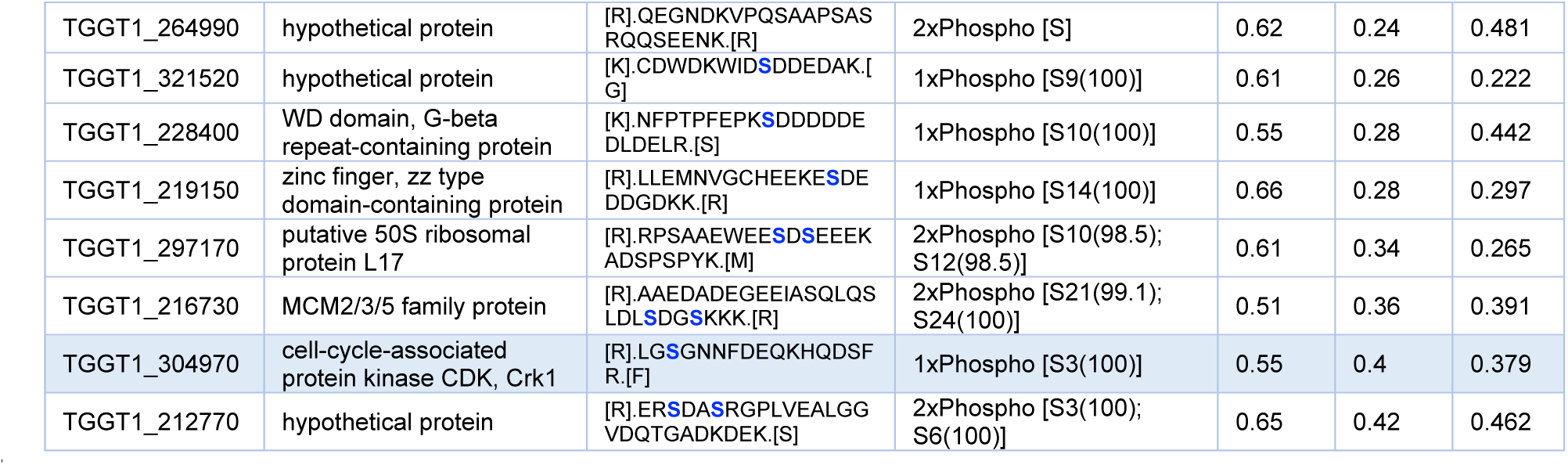
PPKL-dependent phosphopeptides. Listed are proteins with phosphopeptides whose numbers were either increased or decreased in PPKL^AID^ parasites treated with auxin at all time points tested (1, 3, and 6 hours). For the 3 h and 6 h timepoints sample sets, phosphopeptides with FC >= 2 and FC <= 0.5 are listed. For the 1 h timepoint sample set, phosphopeptides with FC >=1.5 and FC <= 0.66 are listed.

Interestingly, among these 29 proteins, two were identified as putative interactors and/or substrates of PPKL via TurboID: SPM1 (TGGT1_263520) and TGGT1_204160 (Fig. 7B, Highlighted in Table 2). SPM1, a protein that stabilizes cortical microtubules in *Toxoplasma* [40, 41], exhibited a phosphorylation increase of greater than two-fold on either S14 alone or both S14 and S18 in all three timepoint sample sets. TGGT1_204160, an eIF2α homolog containing a GYF domain, showed decreased phosphorylation upon PPKL depletion. Remarkably, S1326 of the cell cycle-related kinase Crk1 (TGGT1_304970) showed a significant decrease in phosphorylation at all three time points (Fig. 7B, Highlighted in Table 2). Crk1 has been shown to play an essential role in daughter parasite assembly [42]. Thus, it is plausible that the decreased phosphorylation of S1326 in Crk1 in the absence of PPKL may contribute to the severe defect in the formation of daughter parasites in PPKL-depleted parasites.

## Discussion

In this study, we conducted a comprehensive investigation into the localization and function of PPKL in *Toxoplasma*. Our findings demonstrated that PPKL’s localization is closely associated with the formation of daughter parasites and that depletion of PPKL has a profound impact on the initiation and development of daughter parasites, highlighting the critical regulatory role of PPKL throughout the process of daughter parasite formation.

*Toxoplasma* adopts a unique division process called endodyogeny, where two daughter parasites are generated within the mother cell [34]. The assembly of daughter parasites initiates during the late S-phase of the cell cycle [34, 43]. Each daughter cell is assembled around a centrosome that has already completed the replication process during the early S-phase [34, 43]. The exact composition of the initially assembled daughter parasites is unclear, while it is evident that daughter parasites assemble from the apical to the basal end [34, 43]. Previous studies have identified several IMC proteins, such as IMC15, IMC29, and IMC32, which are present early in the nascent daughter parasites [26, 44, 45]. Additionally, the apical cap proteins ISP1, FBX01, and the AC9-AC10-ERK7 complex, were also identified as the earliest components of daughter parasites [46–48]. However, these previous studies relied on traditional IFA to identify the localization of these proteins, which has limitations in detecting daughter parasites at a very early stage, specifically before the assembly of the IMC. Fortunately, by utilizing U-ExM, we were able to observe newly initiated daughter parasites comprising only two rings: a potentially preconoidal ring labeled by PPKL and a presumed apical polar ring labeled by anti-acetylated tubulin antibody (Fig. 2E). This discovery suggests that PPKL appears in daughter parasites earlier than all the early proteins identified thus far and that the structures of the daughter parasite bud that were first assembled were probably the preconoidal region and the apical polar ring.

The mechanism which regulates the initiation of daughter parasite assembly also remains enigmatic. One of the prominent phenotypes observed upon PPKL depletion in our study was a significant impairment in the initiation of daughter parasites. After an 18-hour culture period, a substantial portion of PPKL-deficient parasites failed to develop detectable daughter parasites. Furthermore, when PPKL-depleted parasites were subjected to a brief 6-hour treatment with IAA, most of the dividing parasites exhibited only a single daughter parasite. These findings indicate that the absence of PPKL disrupts the regulatory pathway responsible for initiating daughter parasite formation. Previous studies have highlighted the requirement of centrosome replication for proper cell division in *Toxoplasma*, with NEK1 and MAPKL identified as protein kinases involved in regulating centrosome duplication [35, 36]. Surprisingly, our investigation revealed that centrosome duplication remained unaffected in PPKL-depleted parasites. Additionally, our phosphoproteomics analysis did not indicate any impact of PPKL depletion on the phosphorylation status of NEK1 and MAPKL. These findings indicate that PPKL potentially participates in a distinct pathway, separate from the one responsible for regulating centrosome replication associated with NEK1 and MAPKL.

Interestingly, our phosphoproteomics analysis revealed that the depletion of PPKL promptly leads to a reduction in the phosphorylation level of Crk1 at S1326, strongly suggesting that PPKL plays a role in regulating the phosphorylation state of S1326 in Crk1. In *Toxoplasma*, Crk1 is an essential cell cycle-associated kinase that partners with the cyclin protein CycL [42]. Although the functional role of the Crk1-CycL complex is not well understood, conditional knockdown of Crk1 results in abnormal assembly of the daughter parasite cytoskeleton [42], suggesting that it is involved in regulating the assembly of the daughter parasite scaffold. In mammals, most cyclin-dependent kinases (CDKs) are known to be activated in a two-step process with the binding of a specific cyclin and the phosphorylation of the activation loop (T-loop) in the kinase domain, typically at a conserved threonine residue [49]. Although the T-loop is present in Crk1, no phosphorylation of this conserved threonine has been reported based on available post-translational modification data of Crk1 (ToxoDB). Notably, S1326, which is ∼30 aa away from the kinase domain, represents the sole identified phosphorylation site within the C-terminal region, encompassing the entire kinase domain (aa 993-1292) and the C-terminal extension (aa 1293-1373), indicating it may be a critical residue for regulating the activity of the kinase domain.

Given that PPKL is a phosphatase and the effect on Crk1 S1326 in its absence is a reduction in phosphorylation, the relation between PPKL and Crk1 is unlikely to be a direct one. The plausibly simplest model is that PPKL activates a kinase that subsequently phosphorylates Crk1. The kinase DYRK1, which we identified as physically interacting with PPKL, could be considered a candidate for such a role. Interestingly, the knockout of DYRK1 in our study causes significantly unsynchronized division of the parasites, highlighting its critical role in regulating parasite division. However, the phosphorylation state of DYRK1 does not have notable alteration (1.23-fold change in 6h samples) with the depletion of PPKL. An alternative model for the functional relation between PPKL and Crk1 is that the phosphorylation of S1326 is the result of autophosphorylation. Under such a model, PPKL would dephosphorylate Crk1 activating its kinase domain and resulting in Crk1 autophosphorylation. Nonetheless, we did not identify Crk1 as a near neighbor of PPKL. Overall, further investigations are necessary to elucidate the precise mechanism of the functional relationship between PPKL and Crk1.

Besides the defect in daughter cell assembly, loss of PPKL resulted in structural changes to cortical microtubules. This finding is consistent with a previous study conducted on *Plasmodium*, which demonstrated that the deletion of PPKL in the parasite led to the dissociation of apical microtubules from the IMC [25]. The cortical microtubules play a crucial role in maintaining the shape and stability of the parasite [41]. The normal cortical microtubule architecture of *Toxoplasma* has robust rigidity and can withstand strong detergent extraction [41]. Our experiments using sodium deoxycholate revealed that PPKL-depleted parasites have more fragile microtubules, suggesting that PPKL is associated with the regulation of the rigidity of cortical microtubules. Along with the loss of rigidity, the cortical microtubules of PPKL-depleted daughter parasites also lose their compact and ordered structure. In those PPKL-depleted parasites undergoing their first cell division, we found that most of those single abnormal daughter parasites had a round shape with much larger gaps between microtubules. This may not only be because these single-daughter parasites are not spatially restricted in the mother parasites without a sibling but also may be mainly due to the loss of the compact and ordered structure of their cortical microtubules. In addition, we also see many other types of irregularly shaped daughter parasites in PPKL-depleted parasites, further supporting PPKL regulates the ordered structure maintenance of the cortical microtubules.

The identification of SPM1 as a putative substrate of PPKL might reveal a mechanism by which PPKL regulates microtubules. SPM1 is a filamentous microtubule inner protein, which binds to α, and β tubulins through the whole length of the microtubule [40, 41]. In addition to SPM1, there are two globular microtubule inner proteins in *Toxoplasma*, TrxL1 and TrxL2, bound to the microtubules [41, 50]. Previous detergent extraction experiments have shown that SPM1 plays a more important role in stabilizing microtubules than both TrxL1 and TrxL2, as microtubules without SPM1 cannot withstand even mild detergent extraction, while microtubules without either TrxL1 or TrxL2 can still withstand strong detergent extraction [41]. Our data suggest that PPKL may regulate the SPM1 by dephosphorylating two residues, S14 and S18. Further studies are needed to verify that changes in the phosphorylation state of these two residues have a significant effect on the stability of the microtubule.

PPKL exhibits an intriguing localization pattern in both mother and daughter cells of *Toxoplasma*, specifically in the preconoidal region. This localization suggests that PPKL may have an undiscovered role in this particular region. Our ongoing research aims to elucidate the specific function of PPKL in the preconoidal region. Furthermore, we have observed that PPKL localizes to the basal complex ring of daughter parasites during the late stages of division, and the basal complex protein BCC7 is one of its neighboring proteins, indicating a potential functional role of PPKL in the basal complex. However, previous proximity-based biotinylation labeling by multiple basal complex proteins did not identify PPKL [51]. Thus it is likely that PPKL is associated with the microtubule ends but not the basal complex itself. Accordingly, its role there may be related to the termination of the extension of microtubules since, as stated above, we have found that many PPKL-depleted parasites have elongated microtubules that run the full length of the parasite.

The closest homolog of PPKL in plants, Bsu1, positively regulates the brassinosteroid signaling pathway through the dephosphorylation of a conserved tyrosine in the CMGC family protein kinase Bin2 [10]. Interestingly, *Toxoplasma* encodes for a Bin2 homolog (TGGT1_265330) that is known to be phosphorylated at the corresponding tyrosine. Nonetheless, we did not identify TgBin2 as either an interactor or a putative substrate in our unbiased approaches. Interestingly, DYRK1, which has been identified as a putative functional partner of PPKL in this study, also belongs to the CMGC kinase family and possesses a phosphorylated tyrosine within a conserved region similar to the phosphorylated tyrosine region of Bin2. This intriguing observation suggests the possibility that PPKL may regulate the activity of DYRK1 by dephosphorylating this conserved tyrosine. Our ongoing research aims to investigate this potential relationship further.

All these data taken together provide a picture of the various roles potentially played by PPKL in *Toxoplasma* and allow us to propose a preliminary model (Fig. 8). We propose that PPKL acts as a key regulator of daughter parasite development in *Toxoplasma*. The dynamic localization of PPKL at different stages of the cell cycle would allow it to precisely regulate the formation and development of daughter parasites as division progresses. The regulation of daughter parasite development by PPKL may begin with the regulation of an unknown pathway that activates Crk1. As division begins, PPKL is recruited earliest formed preconoidal region, although the role it plays there is unknown. With the assembly of microtubules, PPKL appears to play a role in maintaining the rigidity and compact structure of the cortical microtubules, probably by dephosphorylation of microtubule-associated proteins SPM1. Interestingly, at the late stage of daughter parasite development, PPKL enriches the basal complex ring. It is not known what role it plays at this stage, but our data suggest that it may regulate the length of the microtubules, as we observed that in many PPKL-depleted parasites, the microtubules extended to the very bottom of the cell (Fig. 3E bottom).

**Figure 8.**
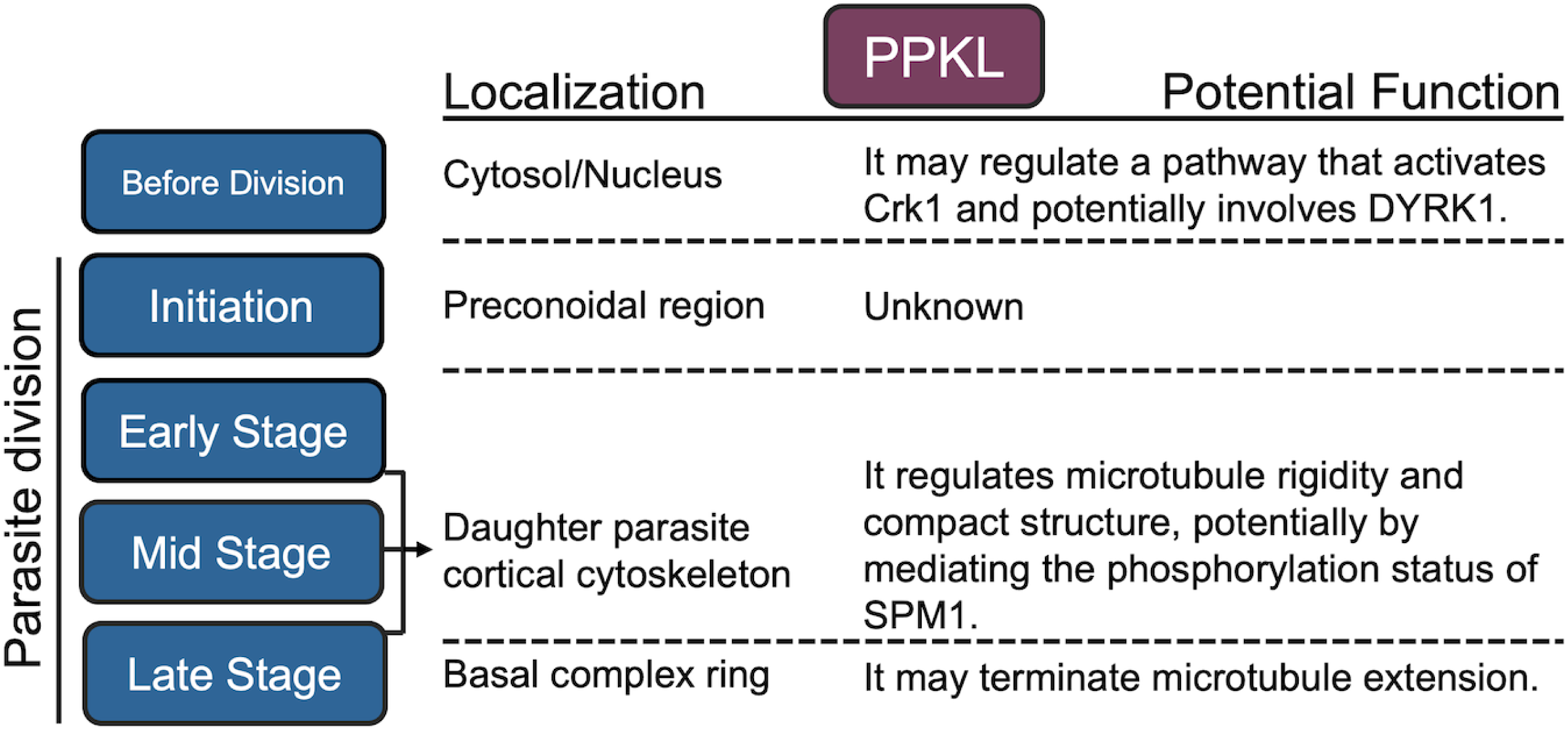
A proposed functional model for PPKL. The localization and potential function of PPKL in the different stages of division (left most column) are listed in the second and third columns.

Overall, the findings of this study highlight the key role of PPKL, a plant-like protein phosphatase, in governing the development processes of daughter parasites in *Toxoplasma*. While potential functional relationships between PPKL and Crk1, as well as PPKL and SPM1, have been identified, the precise molecular mechanisms underlying PPKL’s involvement in the various stages of daughter parasite development remain largely elusive. Further comprehensive investigations into the regulatory mechanisms related to PPKL would greatly contribute to our understanding of the initiation and subsequent development processes of daughter parasites in *Toxoplasma*. Such studies hold great promise in providing exciting prospects for the development of antiparasitic drugs.

## Materials and Methods

### Parasite cultures

*Toxoplasma* tachyzoites used in this study were maintained in human foreskin fibroblasts (HFF) with standard growth medium as previously described [52]. Wild-type parasites used in this study include the strain RH lacking HXGPRT and Ku80 (RH*Δku80Δhxgprt*, referred to as *Δku80*) [53] and the strain stably expressing the plant auxin receptor transport inhibitor response 1 (TIR1) (RH*Δku80Δhxgprt*::TIR1, referred to as TIR1) [33].

### Generation of parasite lines

All primers used for molecular cloning and site-directed mutagenesis are listed in Supplemental Dataset 5. To tag endogenous genes with a hemagglutinin (HA) or Myc epitope tag in the C-terminus, we amplified 3xHA/Myc-DHFR/HXGPRT amplicons from the plasmids LIC-3xHA-DHFR/HXGPRT or LIC-3xMyc-DHFR/HXGPRT with primers containing a 5’ overhang identical to the sequence immediately upstream of the stop codon, and a 3’ overhang identical to the sequence after the Cas9 cutting site. To direct these templates to the desired locus, we generated CRISPR/Cas9 vectors by mutating the UPRT guide RNA sequence in the plasmid pSag1-Cas9-U6-sgUPRT [33] to a guide RNA sequence of the target gene by using Q5 Site-Directed Mutagenesis Kit (NEB). The CRISPR/Cas9 plasmid and the PCR amplicon were transfected into corresponding parental parasites by using the Lonza Nucleofector and Manufacture suggested protocols. Transfected parasites were selected with either pyrimethamine (for selection of DHFR cassette) or MPA/Xanthine (for selection of HXGPRT cassette) and then cloned by limiting dilution as previously described [54].

The PPKL conditional knockdown parasite line was generated based on the same CRISPR/Cas9 mediated strategy described above. The amplicon with regions of homology with the PPKL locus was amplified from the plasmid pAID-3xHA-DHFR [33]. The same CRISPR/Cas9 plasmid used for PPKL endogenous 3’ tagging was transfected together with the AID-3xHA-DHFR amplicon into the TIR1-expressing parasites [33]. DYRK1 knockout parasite strains were generated based on the CRISPR/Cas9 mediated strategy described previously [54]. Briefly, two guide RNA target sites were separately selected in the first exon, and the 3’ UTR of the DYRK1 gene and two CRISPR/Cas9 plasmids were generated. The guide RNA expression cassette from one plasmid was amplified and inserted into the KpnI cutting site of the other plasmid to generate a CRISPR/Cas9 plasmid that expresses two guide RNAs. An amplicon of the DHFR selection cassette was co-transfected into Δ*ku80* parasites together with the double-guide RNA CRISPR/Cas9 plasmid to bridge the two cut sites via homologous recombination.

### Plaque assays

Standard plaque assays were performed as described before [52]. Briefly, 500 parasites of each parasite strain were seeded into host cell monolayers grown in 12-well plates, and cultures were then grown for six days. Cultures were then fixed and stained with crystal violet. Host cell plaques were quantified as previously described [52].

### Immunofluorescence assays

Immunofluorescence assays (IFAs) were performed as previously described [52]. The primary antibodies used include rabbit/mouse anti-HA/Myc (Cell Signaling Technology), rabbit anti-IMC6 (1:1000), rabbit anti-Centrin1 (1:1000), rat anti-IMC3 (1:1000), and rabbit/mouse anti-acetylated-Tubulin (1:5000). A Nikon Eclipse E100080i microscope was used for imaging.

### Ultrastructure Expansion Microscopy (U-ExM)

Ultrastructure Expansion Microscopy (U-ExM) was performed with intracellular and extracellular parasites as previously described [30, 31]. The antibodies used include rabbit anti-HA (1:500) and anti-Centrin1 (1:500), and mouse anti-acetylated-Tubulin (1:500). The fluorescent antibodies used include Alexa Fluor 405 NHS-Ester (1:250), Alexa Fluor 594 (1:500), 488(1:500), and DRAQ5^−^ Fluorescent Probe (1:250). LSM 800 and 900 microscopes were used for imaging using previously described parameters [31].

### Western blots

Western blots were performed as described previously [52]. The primary antibodies used include rabbit anti-HA, anti-eIF2, anti-histone H3, and mouse anti-Sag1. The secondary antibodies utilized were HRP-labeled Anti-Mouse and Anti-Rabbit IgG. The primary antibodies were used at a dilution of 1:5,000, while the secondary antibodies were used at a dilution of 1:10,000.

### Sodium Deoxycholate extraction

Parasite extraction using Sodium Deoxycholate was conducted following the previously described method [55]. Briefly, PPKL^AID^ parasites were cultured in host cells overnight and subsequently treated with either auxin or ETOH for 6 hours. Intracellular parasites were then released using syringe lysis, and the parasites were deposited onto poly-L-lysine-coated coverslips by centrifugation at 100 g for 1 minute. Subsequently, the parasites were exposed to 10 mM Sodium Deoxycholate for 20 minutes at room temperature. Afterward, the parasites were fixed using cold methanol for 8 minutes, followed by IFA utilizing anti-acetylated tubulin and anti-IMC6 antibodies.

### Crosslinking and immunoprecipitation

Crosslinking and immunoprecipitation were performed as previously described [54] with some modifications. Briefly, intracellular parasites (PPKL-3xHA or the parental Δ*ku80* strain) grown in host cells for 24-28 hours were harvested together with host cells by scrapping in cold PBS. After centrifugation, the pellet was resuspended in PBS supplemented with 5mM DSSO (Thermo Scientific) and incubated at room temperature for 10 minutes. Crosslinking was quenched by adding Tris buffer (1 M, pH 8.0) to a final concentration of 20 mM. After two washes with PBS, the samples were lysed with 1 ml RIPA lysis buffer supplemented with protease and phosphatase inhibitor cocktail (Thermo Scientific) at 4°C for 1h. The lysate was then incubated with mouse IgG magnetic beads overnight at 4°C for pre-clearing and then incubated with mouse anti-HA magnetic beads for 4 hours at 4°C. After washes with RIPA lysis buffer and PBS, the beads were submitted to the Indiana University School of Medicine Proteomics Core facility for liquid chromatography coupled to tandem mass spectrometry (LC/MS-MS) analysis.

### Biotinylation by TurboID

PPKL^TurboID-3xHA^ or the control Δ*ku80* parasites were cultured in host cells for around 24 hours. The medium was supplemented with D-biotin (dissolved in DMSO) to a final concentration of 200 μM, and cultures were incubated for 3 hours before harvesting by scraping in cold PBS. The samples were washed with cold PBS three times to eliminate biotin. Then the samples were lysed with 1 ml cold RIPA lysis buffer supplemented with 1x protease inhibitor cocktail (Thermo Scientific) and 1mM PMSF for 1h at 4 °C. After centrifugation, the supernatant of the lysate was incubated with DynaBeads MyOne Streptavidin CI beads (Invitrogen) overnight at 4 °C. The beads were washed two times with 1 ml of RIPA lysis buffer, once with 1 ml of 1M KCl, once with 1 ml of 0.1 M Na_2_CO_3_, and once with 1 ml of 2 M urea in 10 mM Tris-HCl (pH 8.0) and two times with 1ml of RIPA lysis buffer. The beads were lastly washed with PBS two times and submitted to the Indiana University School of Medicine Proteomics Core facility for LC/MS-MS.

### Phylogenetic analysis

The DYRKs used in the phylogenetic analysis including HSDYRK1A (Q13627), HSDYRK1B (Q9Y463), HSDYRK2 (Q92630), HSDYRK3 (O43781), HSDYRK4 (Q9NR20), ATDYRK1 (AT3G177500, ATDYRK2A (AT1G73460), ATDYRK2B (AT1G73450), ATDYRK3 (AT2G40120), TGDYRK1 (TGGT1_204280) and TGDYRK2 (TGGT1_283480). The sequence alignment was performed using the MUSCLE online service, the conserved regions used for tree construction were extracted by using Gblocks 0.91b, and the phylogenetic tree was constructed with a maximum-likelihood method by using PhyML 3.0 with the LG model. The bootstrap values shown on the phylogenetic tree were obtained by repeating the generation of the phylogenetic tree 100 times.

### Phosphoproteomics analysis

Sample preparation, mass spectrometry analysis, bioinformatics, and data evaluation were performed in collaboration with the Center for Proteome Analysis at the Indiana University School of Medicine. Methods described below are adaptations from literature reports [56] and vendor-provided protocols.

Fifteen samples (Experiment 1: n=3 control, IAA 6h; Experiment 2: n=3 control, IAA 1 h, and IAA 3 h) submitted to the Center for Proteome Analysis were denatured in 8 M urea (CHEBI: 16199), 100 mM Tris-HCl, pH 8.5 (CHEBI: 975446756, Sigma-Aldrich Cat No: 10812846001) with sonication using a Bioruptor® sonication system (Diagenode Inc. USA, North America cat number B01020001) with 30 sec/30 sec on/off cycles for 15 minutes in a water bath at 4 °C. After subsequent centrifugation at 14,000g for 20 min, protein concentrations were determined by Bradford protein assay (BioRad Cat No: 5000006). Approximately 2 mg equivalent of protein from each sample was then reduced with 5 mM tris(2-carboxyethyl)phosphine hydrochloride (TCEP, Sigma-Aldrich Cat No: C4706) for 30 minutes at room temperature and alkylated with 10 mM chloroacetamide (CAA, Sigma Aldrich Cat No: C0267) for 30 min at room temperature in the dark. Samples were diluted with 50 mM Tris.HCl, pH 8.5 to a final urea concentration of 2 M for Trypsin/Lys-C based overnight protein digestion at 37 °C (40 µg of protein used for global proteomics and the remainder for phosphoproteomics, 1:70 protease: substrate ratio, Mass Spectrometry grade, Promega Corporation, Cat No: V5072.) Digestions were acidified with trifluoroacetic acid (TFA, 0.5% v/v) and desalted on Sep-Pak® Vac cartridges (50 mg size for global and 100 mg size for phosphopeptides Waters^TM^ Cat No: WAT054955) with a wash of 1 mL 0.1% TFA followed by elution in 70% acetonitrile 0.1% formic acid (FA). Peptide concentrations were checked by Pierce Quantitative colorimetric assay (Cat No: 23275) and confirmed to be consistent.

For phosphoproteomics, each peptide sample (approximately 2 mg) was tapplied to a Pierce High-Select™ TiO2 Phosphopeptide Enrichment Kit (Thermo Fisher Scientific, Cat No: A32993). After preparing spin tips as per the manufacturer’s instructions, each sample was applied to an individual enrichment tip, washed, and eluted as per the manufacturer’s instructions. The phosphopeptide elution was immediately dried. Globa peptidesl and phosphopeptides were each labeled with Tandem Mass Tag (TMT) reagent (manufactures instructions, 0.3 mg per global sample and 0.5 mg per phosphopeptide sample Thermo Fisher Scientific, TMT™ Isobaric Label Reagent Set; Cat No: 90111 Lot XE342654, see Table X below) for two hours at room temperature, quenched with a final concentration v/v of 0.3% hydroxylamine at room temperature for 15 minutes. Labeled peptides were then mixed and dried by speed vacuum.

For high pH basic fractionation, peptides were reconstituted in 0.1% trifluoroacetic acid and fractionated on Sep-Pak® Vac cartridges using methodology and reagents from Pierce™ High pH reversed-phase peptide fractionation kit (8 fractions for global proteomics and 4 for phosphoproteomics skipping every other; Thermo Fisher Cat No: 84868). Samples were run (1/8^th^ of each global and 1/5^th^ of each phosphopeptide fraction) on an EASY-nLC 1200 HPLC system (SCR: 014993, Thermo Fisher Scientific) coupled to Lumos Orbitrap™ mass spectrometer (Thermo Fisher Scientific.) Peptides were separated on a 25 cm EasySpray™ C18 column (2 μm, 100 Å, 75 μm x 25 cm, Thermo Scientific Cat No: ES902A) at 400 nL/min with a gradient of 4-30% with mobile phase B (Mobile phases A: 0.1% FA, water; B: 0.1% FA, 80% Acetonitrile (Thermo Fisher Scientific Cat No: LS122500)) over 160 minutes, 30-80% B over 10 mins; and dropping from 80-10% B over the final 10 min. The mass spectrometer was operated in positive ion mode with a 4 sec cycle time data-dependent acquisition method with advanced peak determination and Easy-IC (internal calibrant) on. Precursor scans (m/z 375-1600) were done with an orbitrap resolution of 120000, RF lens% 30, maximum inject time 105 ms, AGC target of 100% (4e5), MS2 intensity threshold of 2.5e4, MIPS mode, precursor filter of 70% and 0.7 window, including charges of 2 to 7 for fragmentation with 30-sec dynamic exclusion. MS2 scans were performed with a quadrupole isolation window of 0.7 m/z, 37% HCD CE, 50000 resolution, 200% normalized AGC target (1e5), maximum IT of 86 ms, and fixed first mass of 100 m/z.

Raw files were analyzed in Proteome Discover™ 2.5 (Thermo Fisher Scientific) with a database containing *Toxoplasma gondii* GT1 proteins, UniProt reference *Homo sapiens* proteome, plus common contaminants (Total sequences: 79215). Global and phosphoproteomics SEQUEST HT searches were conducted with a maximum number of 3 missed cleavages, precursor mass tolerance of 10 ppm, and a fragment mass tolerance of 0.02 Da. Static modifications used for the search were 1) carbamidomethylation on cysteine (C) residues; 2) TMT label on lysine (K) residues. Dynamic modifications used for the search were TMT label on N-termini of peptides, oxidation of methionines, phosphorylation on serine, threonine or tyrosine, deamidation of asparagine and glutamine, and acetylation, methionine loss or acetylation with methionine loss on protein N-termini. Percolator False Discovery Rate was set to a strict setting of 0.01 and a relaxed setting of 0.05. IMP-pm-RS node was used for all modification site localization scores. Values from both unique and razor peptides were used for quantification. In the consensus workflows, peptides were normalized by total peptide amount with no scaling. Quantification methods utilized isotopic impurity levels available from Thermo Fisher Scientific. Reporter ion quantification was allowed with an S/N threshold of 5 and a co-isolation threshold of 30%. The resulting grouped abundance values for each sample type, abundance ratio (AR) values, and respective p-values (ANOVA) from Proteome Discover™ were exported to Microsoft Excel.

## Acknowledgment

We thank Drs. Peter Bradley, Marc-Jan Gubbels, and Michael Reese for sharing anti-IMC6, IMC3, and *Toxoplasma*-specific anti-acetylated tubulin antibodies, respectively. We extend our gratitude to Dr. Elena Suvorova for her invaluable suggestions for the project. Mass spectrometry was performed by the Indiana University School of Medicine Center for Proteome Analysis. Acquisition of the IUSM Proteomics instrumentation used for this project was provided by the Indiana University Precision Health Initiative. This research was supported by the National Institutes of Health grants R01AI149766, R01DK124067, and R21AI164619 to G.A. The proteomics work was supported, in part, by the Indiana Clinical and Translational Sciences Institute (UL1TR002529) and the Cancer Center Support Grant for the IU Simon Comprehensive Cancer Center (P30CA082709).

## Author Contributions

Chunlin Yang and Gustavo Arrizabalaga conceived and designed the experiments. Emma H. Doud performed phosphoproteomics analysis. Emily Sampson performed endogenous tagging of PPKL. All other experiments and data analysis were performed by Chunlin Yang. The paper was written by Chunlin Yang and Gustavo Arrizabalaga. Emma H. Doud contributed to the manuscript by writing the methods used for phosphoproteomic analysis.

## Competing Interests

The authors declare no competing interests.

## Figure Legends

**Figure S1. PPKL localizes to the nucleus.** A. Western blot of protein samples after cytoplasmic and nuclear fractionation. Anti-HA was used to detect HA-tagged PPKL. Anti-eIF2α and anti-histoneH3 were used as controls to detect eIF2α, a cytoplasmic protein, and histone H3, a nuclear protein. B. ImageJ was used to quantify the relative intensity of the bands in the two portions labeled by the same antibody. Fisher exact test was used to compare the ratios of cytoplasmic/nuclear of PPKL was significantly different from that of the control eIF2α.

**Figure S2. Fusion of AID to the C-terminus of PPKL reduced its expression.** A. Western blot of protein samples isolated from PPKL^HA^ and PPKL^AID^ parasites. Anti-HA was used to detect PPKL-3xHA and PPKL-AID-3xHA. The protein Sag1 was used as a loading control. B. The quantification of the Western blot in panel A reveals the relative expression levels of PPKL-AID-3xHA normalized to Sag1 and PPKL-3xHA.

**Figure S3. PPKL-TurboID validation.** A. Localization of PPKL-TurboID-3xHA in intracellular parasites assessed by IFA. B. Western blot showing biotinylated proteins extracted from PPKL^TurboID^ parasites treated with or without D-biotin. Detection was achieved using Streptavidin-Conjugated Horseradish Peroxidase.

## Supplementary datasets

**Supplementary Dataset 1.** Proteins immunoprecipitated with TgPPKL and identified by LC-MS/MS. The cutoff of fold change is PPKL.3xHA /Control >= 2.

**Supplementary Dataset 2.** List of proteins biotinylated by the PPKL-TurboID fusion. For each repeat, the fold change cutoff was PPKL-TurboID/Control >=2. The list of PPKL neighboring proteins was selected based on the following criteria: in combination with two replicates, 1) identified in both replicates; 2) a total of 10 or more peptides were identified between the two replicates; 2) the fold change was equal to or larger than 3.5.

**Supplementary Dataset 3.** Listed are phosphopeptides identified in PPKL^AID^ parasites treated with auxin or ethanol for 6 h by phosphoproteomics analysis. The sheet “PeptideGroups” contains all phospho-peptides identified in parasites and host cells. *Toxoplasma* phosphopeptides that were significantly (p-value <= 0.05) increased or decreased in auxin-treated parasites are listed in the sheets titled “Toxo Increased” and “Toxo Decreased”. The phosphopeptides that were increased or decreased by more than two-fold in phosphorylation are listed in “6h Increased FC >2” and “6h Decreased FC < 0.5”. The phosphopeptides that were from the proteins identified by TurboID analysis are listed in “Increase overlap with TurboID” and “Decrease overlap with TurboID”. Proteins in Fig. 8A have been listed in the sheet “Proteins of Fig. 8A”. Those proteins that are PPKL neighboring proteins identified by TurboID analysis were highlighted.

**Supplementary Dataset 4.** Listed are phosphopeptides identified in PPKL^AID^ parasites treated with auxin for 1 and 3 h or ethanol for 1 h. The sheet ‘PeptideGroups’ lists all phosphopeptides identified in parasites and host cells. The phosphopeptides identified in parasites are shown in the sheet “Toxo peptides”. The phosphopeptides that are more/less abundant in 1 or 3 h auxin-treated parasites were filtered via specific fold changes and are shown in corresponding sheets. Proteins of Fig. 8A have listed in the sheet “Proteins of Fig. 8A”. Proteins identified as putative PPKL neighboring proteins by TurboID are highlighted.

**Supplementary Dataset 5.** List of primers used in this study.

